# Modeling of GyrA, MexB, FtsI, AtpD Protein Variants In Multidrug Resistant *Acinetobacter baumannii*

**DOI:** 10.1101/2025.01.03.631236

**Authors:** Nik Zakuan Hakim Nik Mohd Nazri, Siti Azma Jusoh, Kirnpal Kaur Banga Singh, Mohamad Izwan Ismail

## Abstract

*Acinetobacter baumannii* is a Gram-negative nosocomial pathogen known to manifest numerous drug resistances against major antibiotic classes. Compounded by its pathogenicity and virulence, it is considered globally as a top priority threat among the ESKAPE pathogens. The GyrA, MexB, FtsI and AtpD proteins in *A. baumannii* strain PR07 have been proven to mutate under exposure to ciprofloxacin, meropenem, imipenem and erythromycin, respectively. While the genomic data is useful, the impact of the mutations on the protein structure and function is not well understood. To obtain a deeper understanding, the protein structures were analyzed using structural bioinformatics tools. Here, the PR07 GyrA, MexB, FtsI and AtpD protein sequence data from NCBI were compared between ESKAPE pathogens and other *A. baumannii* species. MexB and AtpD structures were retrieved from the PDB database, while AlphaFold was used to construct protein structure predictions for GyrA and FtsI. MSA analyses identified mutations GyrA S81L, MexB S181L, FtsI P508, FtsI A515V, FtsI A579T and AtpD A166V mutations to be unique among the selected bacteria species. The mutation sites for all four target proteins were found to be within proximity to the potential binding sites. GyrA S81L, MexB S181L, FtsI A515V and FtsI A579T rigid protein models have shown loss of inter-residue polar hydrogen bonds, while AtpD A166V caused no observable changes. The mutations reported in PR07 therefore may potentially be significant contributors to its acquired resistance towards the target antibiotics.

## 1.0 INTRODUCTION

Antimicrobial resistance (AMR) is a state when bacteria become unaffected by antibiotics, which complicates treatment efforts (Muteeb et al., 2023). Throughout the years, AMR has steadily developed into a global crisis, and the rampant misuse of antibiotics during the COVID-19 pandemic has further worsened the situation (Lai et al., 2021). In 2019, analysis by Muray et al. estimated 1.27 million deaths that were attributable to bacterial AMR out of 4.95 million deaths associated with AMR (Murray et al., 2022).

*Acinetobacter baumannii* is a Gram-negative gammaproteobacteria that is commonly occurring in nosocomial settings, especially in ventilator-associated infections. Due to its genomic plasticity, it has been reported to be capable of relatively rapid mutations and high adaptability towards selective pressure (Baquero et al., 2021). *A. baumannii* has been identified as one of the main causes of hospital-associated infections (HAIs) worldwide. Recent findings have shown the increasing emergence of multidrug resistance (MDR) by the species within a few decades (Alrahmany et al., 2022).

To date, the study of AMR evolution in bacteria remains relatively untapped. While pre-post antibiotics exposure studies are common, detailed insight into the timeline of evolution is not as widely explored. This is likely due to the exponentially higher cost, time commitment and effort needed to complete. Existing studies have suggested that pathogens require a cascade of mutations or other conditions in order to unlock further genomic adaptations (Ismail, 2017). Furthermore, genomics studies alone do not highlight the impact of mutations towards potential protein-drug interactions (Yeh et al., 2009). This is where protein structure studies are essential.

The multidrug efflux pump component MexB protein belongs to a group of RND-type multidrug efflux pumps. It is a homologue to the AdeJ protein and typically forms the MexAB-OprM complex. Mutations in MexB efflux proteins for example have been reported to alter substrate selectivity in *Pseudomonas aeruginosa* (Blair et al., 2015). In addition, modifications in drug-binding pockets have also been suggested to result in clinically relevant resistance (Middlemiss & Poole, 2004).

Mutations in gyrase A (GyrA) have been studied extensively. However, the impact of these mutations on protein structure and function, especially in *A. baumannii* is not well studied. Recent advances in computational analyses revealed that mutations in GyrA are relevant to ciprofloxacin resistance (Singh et al., 2021).

The *ftsI* gene encodes the penicillin-binding protein 3 (PBP3) and is part of a group of genes; *ftsA, ftsEX, ftsK, ftsQ, ftsL, ygbQ, ftsW, ftsI, ftsN*, and *ftsZ* (Wissel & Weiss, 2004). Mutations in this gene have been associated with increased resistance towards beta-lactam antibiotics (Takahata et al., 2007; Witherden et al., 2014) due to its role in cytokinesis and the formation of the bacterial septa (Li et al., 2022; Errington et al., 2003).

The *atpD* gene is a highly conserved region encoding for the ATP synthase F1 subunit β and may be associated with metabolism and biofilm production (Algammal et al., 2021; Colquhoun & Rather, 2020; Rumbo-Feal et al., 2013). Mutations in the ATP synthase have been reported to decrease fitness and are outcompeted by wild-type strains

(Lapashina & Feniouk, 2019). Such metabolic adaptations are suggested to be associated with antimicrobial resistance (Zheng et al., 2022; Ismail, 2017).

Despite being a significant model for antimicrobial resistance evolution, experimental structures for *A. baumannii* associated resistance proteins are limited. This explains the scarcity of studies focusing on structure modification analyses caused by mutations. Furthermore, current *in silico* prediction methods utilizing protein sequences that scrutinize mutation sites rarely include in-depth protein structure analyses, leaving a huge gap in understanding the impact of the mutation on resistance. With the advent of recent protein structure prediction such as AlphaFold (ref), it is now possible to further extrapolate genomic data and elucidate previously untapped structural data.

The previous study by Ismail has successfully identified several mutations suspected to be the key in the resistance development of *A. baumannii* (strain PR07) towards ciprofloxacin, erythromycin, meropenem and imipenem. With recent updates in protein databases and modeling software, we further analyzed these mutations and extrapolated their implications towards protein structure and function. The objectives of this study are therefore to determine; i) the position of the mutation sites within the associated protein structures and ii) the effects of the mutations on the protein structures.

## 2.0 METHOD

### 2.1 Target Protein Selection and Sequence Retrieval

In this study, mutant proteins from *A. baumannii* strain PR07 studied by Ismail (2017) were selected for structural analysis. Initial FASTA sequences were obtained from the *A. baumannii* PR07 genome (accession no: SAMN02471432). The nucleotide sequences were BLASTed against the NCBI database to verify the continued relevance of reported gene nomenclature and function. Subsequently, the presence of suitable reference structures in the UniProt database were verified (Table 1).

**Table 1.** List of *A*.*baumanii* target proteins used in this study. The protein information was retrieved from the UniProt database.

| UniProt ID | Gene | Encoded Protein | AA length | PDB ID |
| --- | --- | --- | --- | --- |
| X2L5Z9 | *gyrA* | DNA gyrase subunit A | 904 | N/A |
| Q2FD94 | *mexB* | Efflux pump membrane transporter MexB | 1058 | 7M4Q |
| G1C6Z8 | *ftsI* | Peptidoglycan D/D-transpeptidase | 610 | 3UE3 |
| V5VHQ6 | *atpD* | ATP synthase subunit beta | 464 | 7P2Y |

### 2.2 Intra- and Interspecies Protein Sequence Analysis

The nucleotide sequences of the target proteins were obtained and BLASTed against the NCBI database. Five hits within the same species were then randomly selected and aligned using ClustalO to verify the PR07 mutant residue (Madeira et al., 2022). The multiple sequence alignment (MSA) was repeated against proteins in ESKAPE pathogens to verify its uniqueness.

### 2.3 Acquisition of Protein 3D Models

The MexB, FtsI and AtpDr *A. baumannii* protein structures were retrieved from the PDB database (**Table 1**). For GyrA, which lacks an *A. baumannii* experimental structure, the 3D model was generated from the AlphaFold database. The quality of the GyrA AlphaFold model was evaluated by Ramachandran plot using the ZLab website (Santra et al., 2022). The heteromultimeric form of the MexB, FtsI and AtpD were directly obtained from the PDB database while FtsWI was constructed using ChimeraX (FtsI AFID: A0A427X5Y8, FtsW AFID: A0A009PYL1) (Meng et al., 2023). To predict GyrAB structure, the AlphaFold structure of *A. baumannii* GyrA and GyrB (GyrA AFID: A0A5K1MHS3, GyrB AFID: F0KPH7) were structurally aligned to the *E. coli* DNA Gyrase.

### 2.4 Generation of Mutant Protein Models

Pymol was employed to prepare the mutant structures. The substitution tool was utilized to replace a mutant residue. The lowest energy side chains were selected for the mutant models. The active sites were predicted using DoGSitescorer, where binding sites with a druggable score ≥ 0.7 were recorded. The internal binding sites with groove shapes are prioritized for presentation purposes. All programs utilized default settings unless specified.

### 2.5 Generation of FtsWI Multimer

To confirm the positioning of the mutant FtsI protein in *A. baumannii*, the multimeric form of FtsWI was constructed using ChimeraX and Google Collab GPU. The PR07 FtsW and FtsI amino acid sequences were used as the input, in which the sequences were comma-separated. The best multimer structure indicated by the Predicted Local Distance Difference (pLDDT) plot and the Predicted Aligned Error (PAE) plot was recorded for further analysis. The overlay of *A. baumannii* FtsWI (Alphafold multimer prediction) with *P. aeruginosa* FtsWI (experimental) was generated using PyMol to compare the contact point/interaction between FtsW and FtsI.

## 3.0 RESULTS

### 3.1 DNA gyrase subunit A, GyrA

The targeted PR07 GyrA protein was confirmed to be 904 amino acids in length, with a Ser81Leu mutation. The intraspecies MSA analysis shows nine of the 11 *A. baumannii* reference strains exhibited a serine residue at position 81, while the other two exhibited a leucine residue at the same position (**Figure 1**). When compared to the eight ESKAPE pathogens, the same serine residue was detected at position 81, with the exception of *S. aureus, K. pneumoniae* and *P. aeruginosa* exhibiting leucine, isoleucine and threonine variants, respectively.

**Figure 1.**
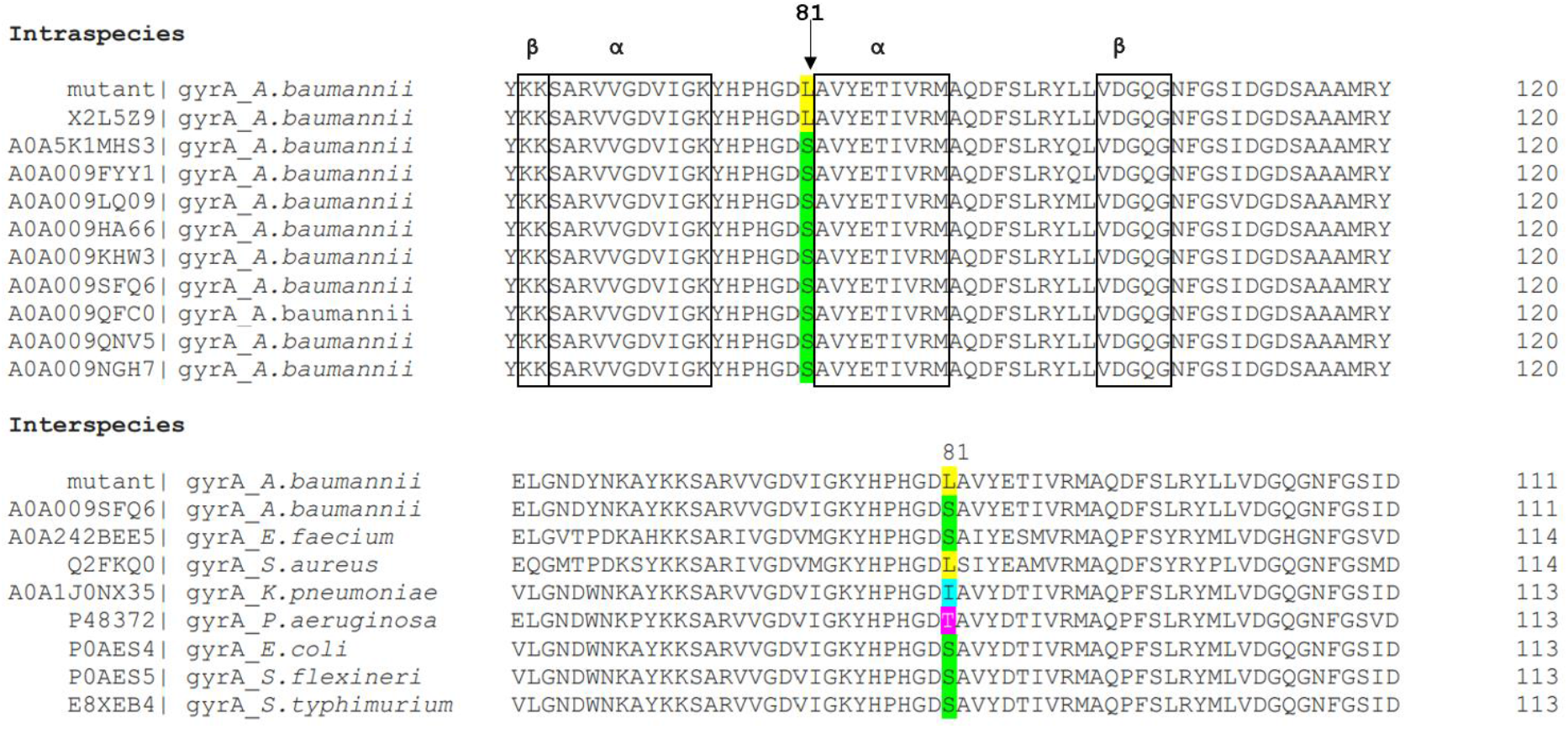
MSA of GyrA protein sequence aligned with *A. baumannii* and selected ESKAPE pathogens. Ser81Leu amino acid variation (highlighted in yellow) is uncommon in *A. baumannii* wild-type strains. The interspecies alignment showed four variations that existed across other ESKAPE pathogens. The Ser81Leu variation was observed in the ciprofloxacin-resistant *A*.*baumannii* culture. The blue boxes represent either alpha or beta-secondary protein structure predictions based on the amino acid sequence.

In order to evaluate the impact of the Ser81Leu mutation in GyrA, a 3D model of the *A. baumannii* protein was acquired from the AlphaFold database, and consists of two GyrA chains and two GyrB chains (**Figure 2A**). The GyrA Ser81Leu mutation site is positioned at the surface of the GyrA where the two GyrA monomers are within proximity to one another, deep in the tetramer complex (**Figure 2B** and **Figure 2C**). Consequently, the mutant residue may cause disruption to the interactions between the monomer subunits. As shown in **Figure 1**, amino acid sequence alignment and secondary structure prediction using Clustal Omega and SWISS-MODEL respectively, revealed that the Ser81Leu mutation (occurring close to the alpha helix of 82-90th amino acids) is comparable to the *E. coli* GyrA Ser83Leu variant (PDBID: 6RKU). In addition, analysis using DogSiteScorer indicated that the Ser81Leu mutation occurs within proximity to at least two predicted druggable binding sites (**Figure A1**).

**Figure 2.**
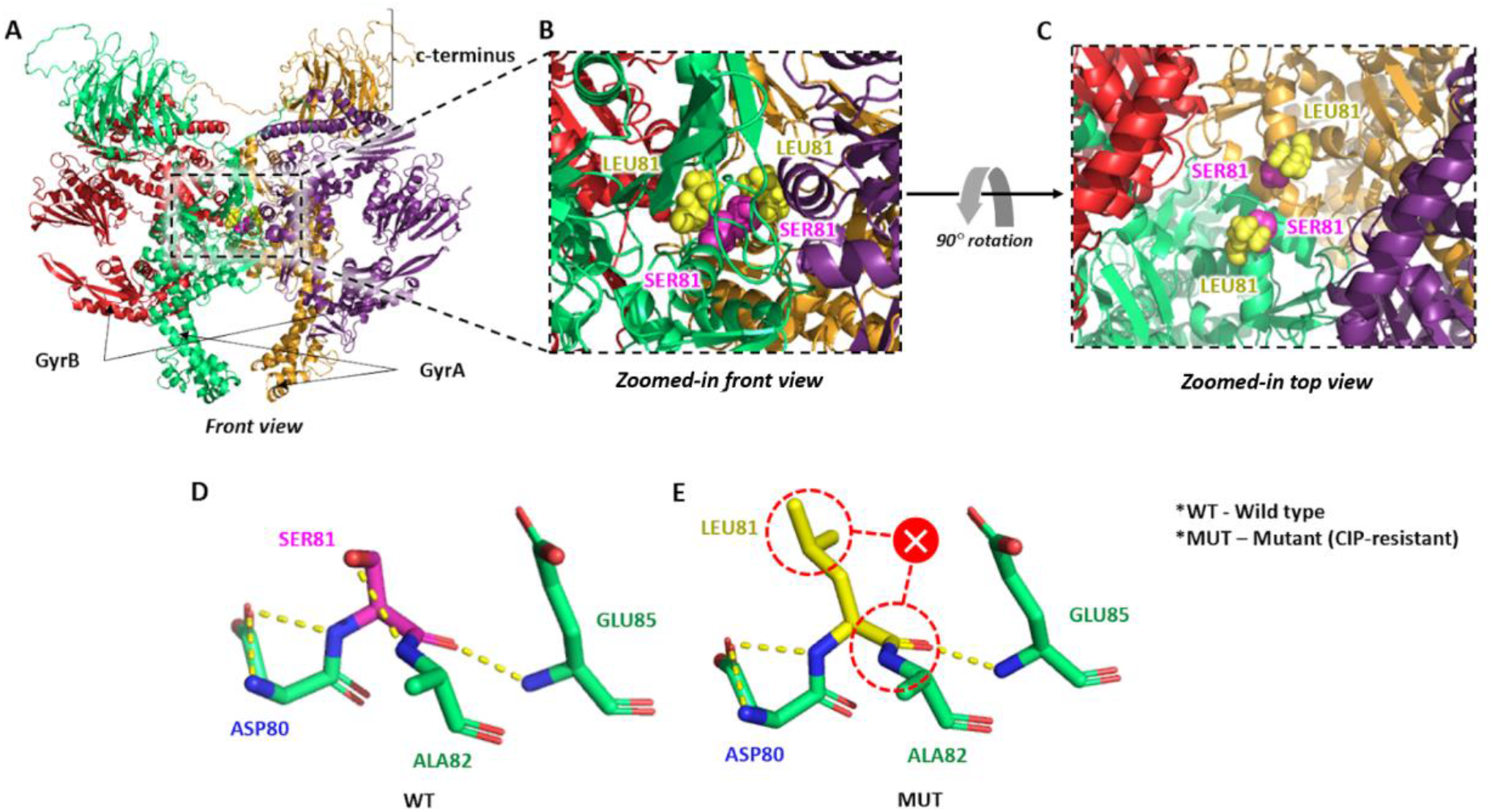
Model structure of *A. baumannii* GyrA predicted by AlphaFold. **A**. The assembly unit of the GyrA heterotetramer consists of two GyrA chains and two GyrB chains. (GyrA AFID: A0A5K1MHS3, GyrB AFID: F0KPH7). **B**. Zoomed-in image of the front view of GyrA focusing on the mutation sites. **C**. Top view of the complex focusing on the mutation sites. Wild type (Ser81) and mutant (Leu81) residues are represented by magenta and yellow spheres, respectively. **D**. Ser81 wild-type residue (WT) has three hydrogen bonds with the neighboring residue **E**. Mutant Leu81 shows a loss of hydrogen bonds with Alanine 82 (denoted by the red X symbol), as shown in the ciprofloxacin-resistant variant (MUT).

Based on the mutagenesis analysis, the Ser81Leu substitution caused a loss of hydrogen bonds between the alcohol functional group in Ser81 and nitrogen in Ala82. Interactions with other surrounding residues, such as hydrogen bonds with Asp80/C, Ala82/N Glu85/N(H) remained unchanged (**Figure 2D** and **Figure 2E**). In addition, Ser81Leu altered the biochemical property of the residue location, which is from an uncharged polar to a hydrophobic side chain. Furthermore, serine is noted to take up 13 atoms of space compared to the mutant leucine, which takes up 22 atoms of space.

### 3.2 Efflux pump membrane transporter, MexB

The MexB monomer from *A. baumannii* PR07 was determined to be composed of 1058 amino acid residues with a Ser181Leu mutation. MSA analysis indicated that the Leu181 mutation is unique to PR07 when compared to the 10 selected *A. baumannii* strain, whereby alanine and glycine were detected instead (**Figure 3**).

**Figure 3.**
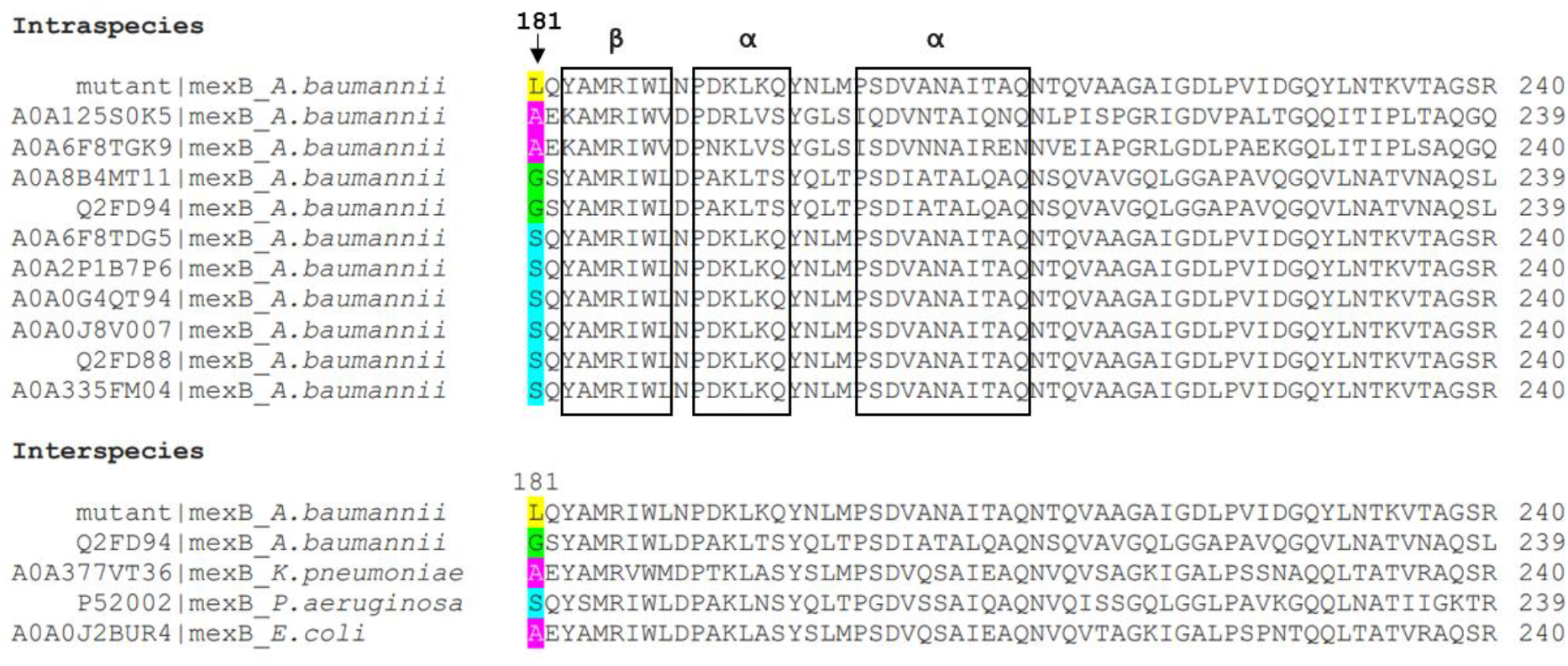
MSA of the mutant PR07 MexB protein with *A. baumannii* and selected ESKAPE pathogens showing Ser181Leu substitution to be uncommon in both alignments. The boxes represent either alpha or beta-secondary protein structure predictions based on the amino acid sequence.

A similar outcome was determined when comparing MexB protein sequences from the ESKAPE pathogens, where alanine and glycine residues were noted at the 181th position in MexB.

Currently, the *A. baumannii* MexAB-OprM complex structure is not available in the PDB database. Therefore, to determine the structural impact, the MexB homotrimer of *A. baumannii* (PDB ID:7M4Q) (**Figure 4**) was superimposed with the MexAB-OprM structure from *P. aeruginosa* efflux pump (PDB ID: 6TA5). While the Ser181Leu mutation did not occur in any of the identified ligand binding sites, it is still within proximity of several druggable pockets (**Figure A2**). It was also noted that the mutation occurs in a primary structure connecting two beta-sheets, between 172-178 (not shown) and 183-189 residues (**Figure 3**). Despite the Ser181Leu residue being located deep within the protein complex, it is positioned on the surface of the internal passageway where molecules pass through the MexB-OprM complex. However, this mutation presumptively does not cause a structural impact on the MexA that is docked above the MexB protein.

**Figure 4.**
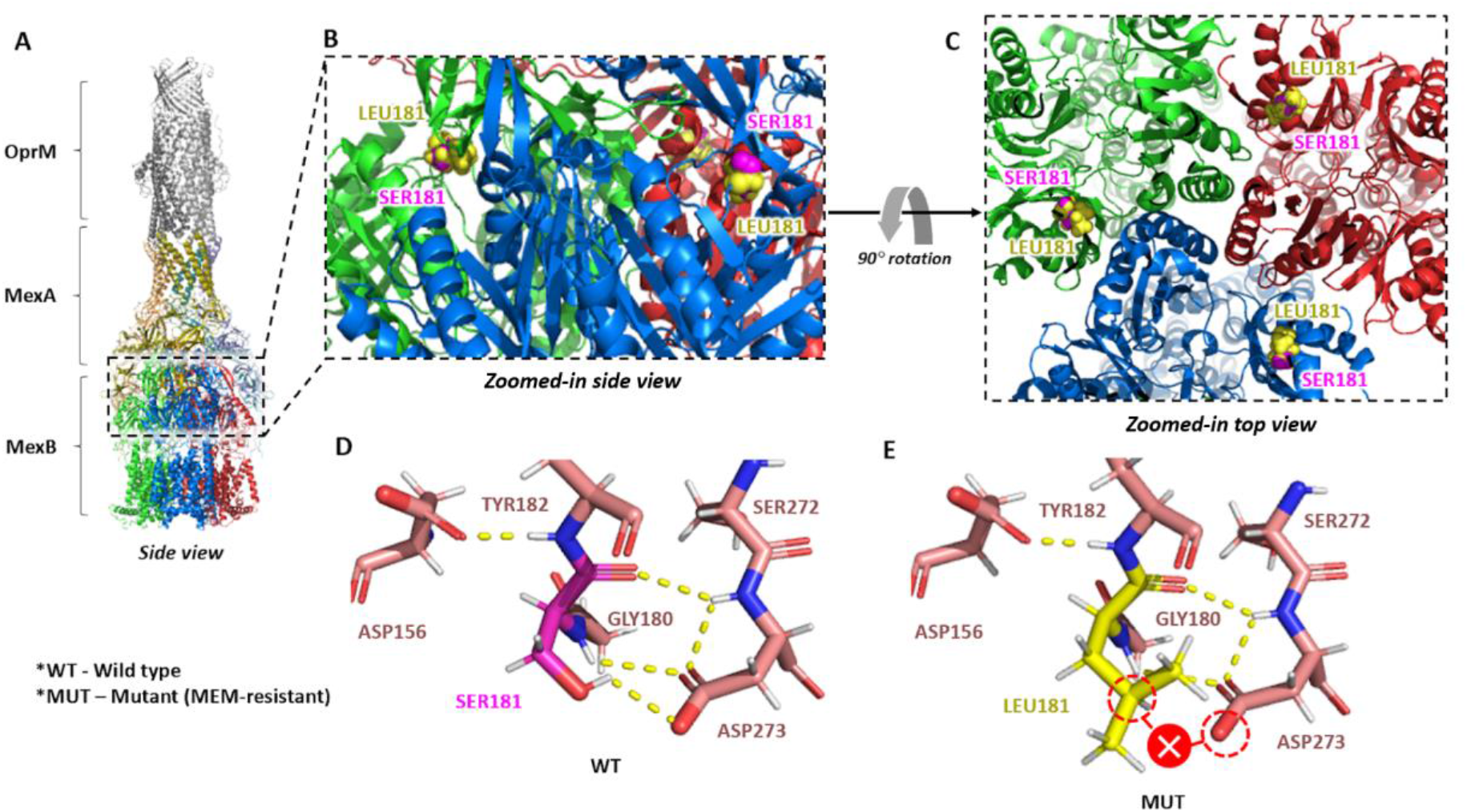
3D model of *A. baumannii* efflux pump membrane transporter, MexB. **A**. Superimposed model of *A. baumannii* MexB homotrimer (PDBID:7M4Q) and *P. aeruginosa* MexAB-OprM protein complex consisting of three OprM chains, six MexA chains and three MexB chains (PDBID: 6TA5). **B**. Side view (zoomed-in) of MexB. **C**. Top view (zoomed-in) of the *A. baumannii* MexB homotrimer showing the wild type, Ser181 and mutant, Leu181 residues represented by magenta and yellow spheres respectively. **D**. PR07 MexB wild-type residue showing Ser181 exhibiting four hydrogen bonds with adjacent residues. **E**. A loss of hydrogen bond between Ser181Leu with Asp273 (indicated by the red X symbol) was predicted in the meropenem-resistant PR07 variant.

Analysis of the residue interactions indicated that the MexB Ser181Leu structure displayed a missing hydrogen bond between Ser181 and Asp273. The peptide bonds with the neighboring Gly180 and Tyr182 and hydrogen bonds with nearby Asp273 remain unchanged. Furthermore, the Ser181Leu resulted in altered amino acid properties (polar uncharged to hydrophobic side chain), as well as a modified spatial occupancy; serine (13 atoms) to leucine (22 atoms).

### 3.3 Peptidoglycan D, FtsI

*A. baumannii* FtsI was confirmed to be 610 amino acids in length. According to the study by Ismail, exposure to imipenem has led to two mutations in the FtsI monomer; Ala515Val and Ala579Thr, while exposure to erythromycin resulted in a single Pro508Leu mutation in a separate lineage. Interspecies MSA of 10 selected *A. baumannii* FtsI proteins indicated Ala515, Ala579 and Pro508 to be the prevalent amino acids in wild-type strains. Interspecies MSA with ESKAPE pathogens on the other hand indicated Leu and Tyr to be prevalent at position 515, Leu and Glu at position 579 and Lys and Ala at position 508 (**Figure 5**).

**Figure 5.**
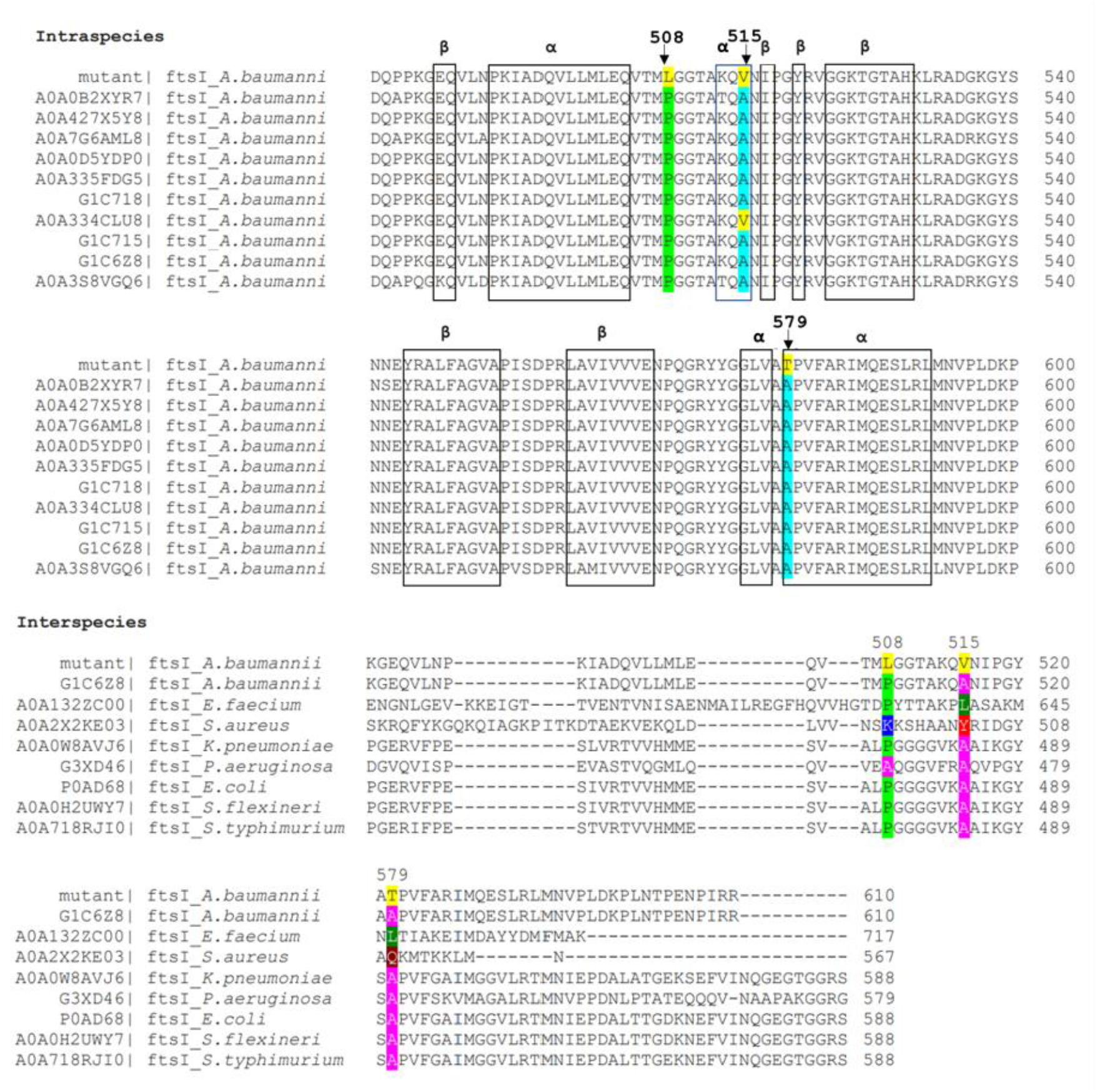
Multiple sequence alignments of FtsI protein sequence with *A. baumannii* and selected ESKAPE pathogens. The Pro508Leu (highlighted in yellow) variation was not found in either the 10 *A. baumannii* wild-type strains or ESKAPE pathogens. Ala515Val and Ala579Thr (both highlighted in yellow) are also uncommon variants when compared to both the intraspecies and interspecies targets. Pro508Leu was determined to be unique to the erythromycin-resistant PR07 strain, while both Ala515Val and Ala579Thr mutations are unique to the imipenem-resistant PR07 strain. The blue boxes represent either alpha or beta-secondary protein structure predictions based on the amino acid sequence.

The available *A. baumannii* FtsI experimental structure (PDBID: 3UE3) did not cover the transmembrane portion (1-64 aa position, alpha helix structure) of the PR07 FtsI. Therefore, the FtsI AF structure was hetero-dimerized with FtsW to construct *A. baumannii* FtsWI via AF multimer prediction. The FtsI transmembrane portion is essential as it interacts with and allows the dimerization with FtsW, similarly with *P. aeruginosa* FtsWIQBL complex (PDB ID: 8BH1). Furthermore, superimposition of the predicted FtsWI heterodimer and *P. aeruginosa* FtsWIQBL is in consensus with the contact points of ftsI and ftsW, which are FtsI transmembrane (TM) with TM8 and TM9 of FtsW, as mentioned by Kashammer et al., 2023.

Ala515Val and Ala579Thr mutations in FtsI are located within proximity to the C-terminus of the polypeptide chain of the transmembrane region of the protein (**Figure 5**), while Pro508Leu occurs in between two alpha-helices (492-504 and 513-515). Alternatively, both Ala515Val and Ala579Thr occur within an alpha helix, with the former being at the end of a three-amino acid alpha helix (513-515), while the latter is at the start of a 14-aa alpha-helix (579-592) (**Figure 6B**). All three mutations occurred close to the known binding sites as predicted by DogSiteScorer **(Figure A4**).

**Figure 6.**
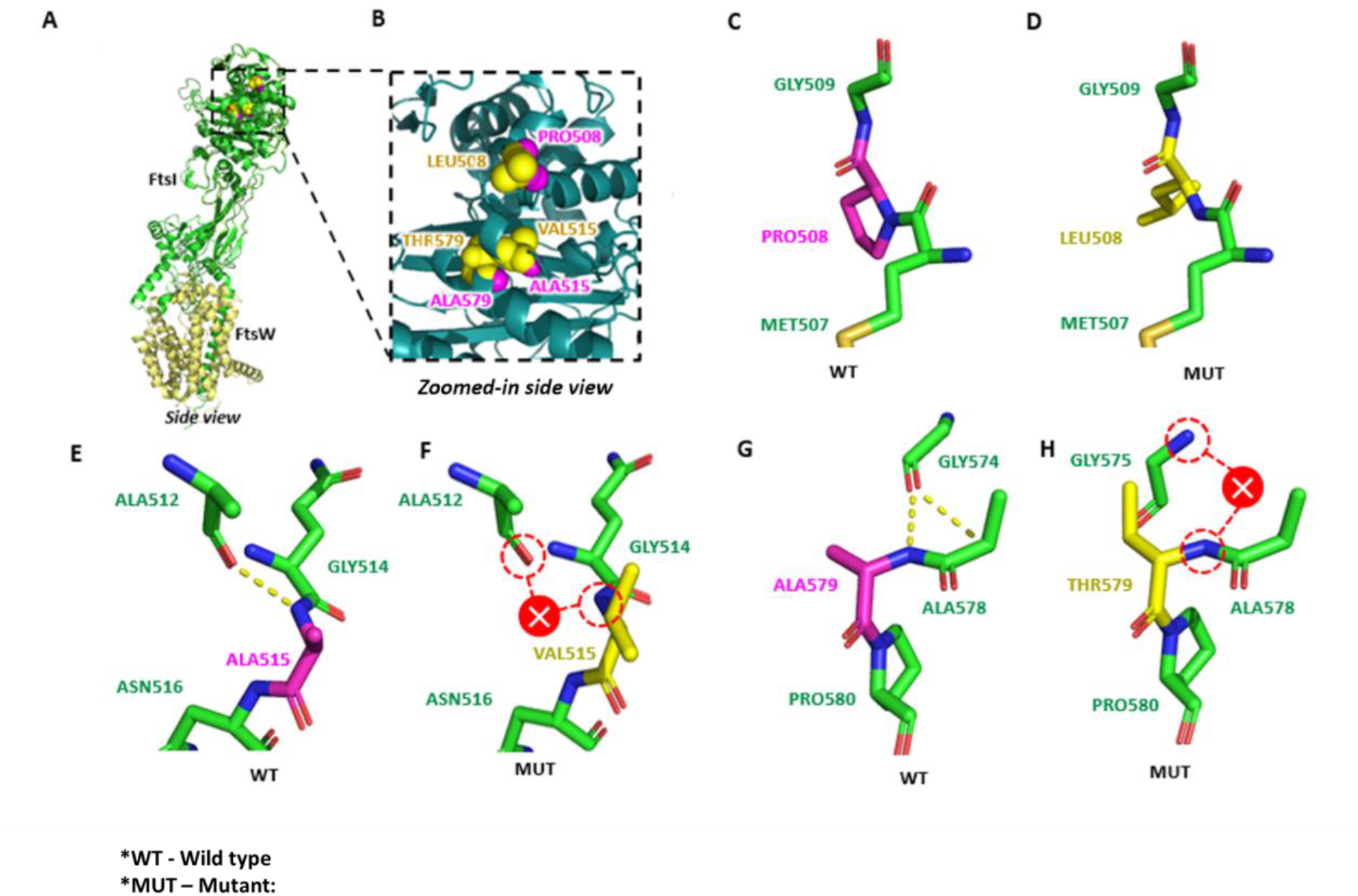
Model structure and mutation sites of *A. baumannii* transpeptidase, FtsI. **A**. Side view of predicted FtsWI heterodimer from A. baumannii PR07 strain. It consists of a single FtsI chain (AlphaFold ID: A0A427X5Y8) and a FtsW chain (FtsW AlphaFold ID: A0A009PYL1). **B**. Zoomed-in image of the *A. baumannii* FtsI region displaying three mutation sites combined from two separate lineages; Erythromycin-resistant strain (Pro508Leu), and meropenem-resistant strain (Ala515Val and Ala579Thr). The wild-type (WT) and mutant (MUT) residues are represented by magenta and yellow spheres respectively. **C**. Pro508 wildtype residue. **D**. Leu50 mutant residue8. **E**. Wild-type Ala515 forms a hydrogen bond with Ala512. **F**. Mutant Val515 caused a loss of hydrogen bond with Ala512. **G**. Wild-type Ala579 forms a hydrogen bond with Gly574. **H**. Mutant Thr579 caused a loss of hydrogen bond with Gly574.

Substitution of Pro508Leu in the FtsI protein model did not result in any observable changes to the adjacent amino acids (**Figure 6C** and **Figure 6D**). Ala515Val however was noted to lose its hydrogen bond with a nearby Ala512, while Ala579Thr lost its hydrogen bond with Gly574 (**Figure 6E-H)**. Amino acid property changes were identified in Pro508Leu (nonpolar to hydrophobic) and Ala579Thr (hydrophobic to non-polar), as opposed to Ala515Val (hydrophobic). The spatial occupancy of the mutation sites were noticeably altered; Pro508Leu (17 atoms to 22 atoms), Ala515Val (13 atoms to 19 atoms), and Ala579Thr (13 atoms to 17 atoms).

### 3.4 ATP synthase subunit beta, AtpD

The *A. baumannii atpD* gene encodes the ATP synthase subunit beta protein (AtpD) which consists of 464 amino acid residues. The study by Ismail reported that prolonged exposure to meropenem resulted in the Ala166Val mutation in this subunit. Both the inter- and intraspecies MSA analyses revealed the mutation to be unique to the PR07 variant (**Figure 7**).

**Figure 7.**
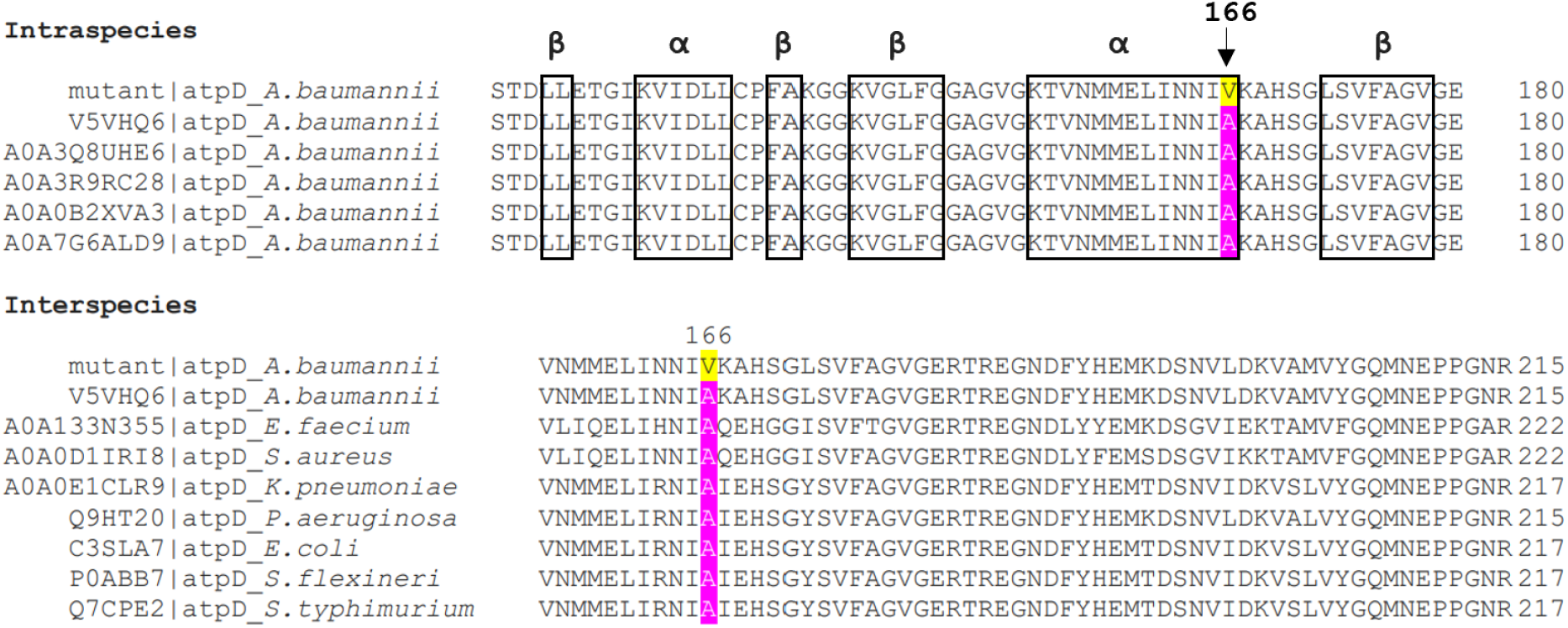
Inter- and intraspecies MSA of the AtpD protein indicates the Ala166Val mutation to be unique to the meropenem-resistant PR07 strain. The blue boxes represent either alpha or beta-secondary protein structure predictions based on the amino acid sequence.

As shown in **Figure 7**, the Ala166Val mutation occurred at the end of the α-helix structure (154-166). Structural analysis of the mutation site indicates that the mutation is located at the outward-facing surface of the AtpD (**Figure 8B**). The location of the substituted Val166 is distal from the neighboring AtpA subunits on either side, which resulted in no apparent cross-subunit interactions. Thus, the formation of AtpD-AtpA heterohexamer may not have been disrupted by the residue substitution.

**Figure 8.**
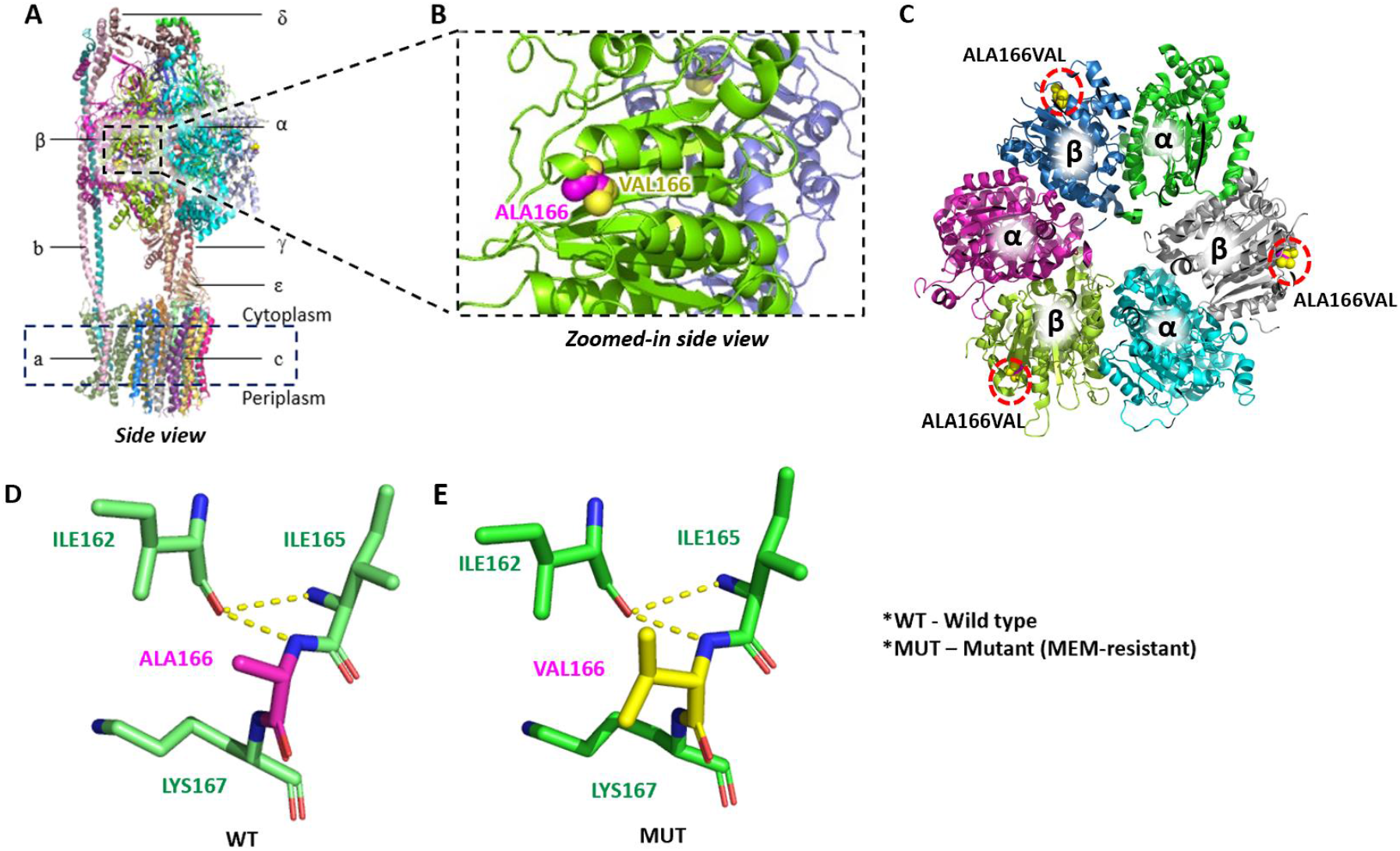
Structure of the *A. baumannii* ATP synthase subunit beta, AtpD (PDBID: 7P2Y) **A**. The functional assembly of *A*.*baumannii*, consisting of α, β, γ, δ, ε, a, b and c chains. **B**. Zoomed-in side view of a single AtpD monomer showing the wild type residue, Ala166 and the mutant residue, Val166 represented by magenta and yellow spheres, respectively. **C**. Ala166Val was located at the outer surface of AtpD (viewed from the top). AtpD is labeled as β-chain while AtpA is labeled as α-chain **D**. Wild type residue, Ala166 exhibits hydrogen bond interactions with a nearby Ile165 **E**. Mutant residue, Val166 also exhibits the same interactions as Ala166..

As shown in **Figure 8**, the structural analysis of the amino acid substitution of AtpD indicates no apparent changes when Ala166 is replaced by Val166. The mutant amino acid exhibits similar physicochemical properties as the wild type residue (small hydrophobic).

## 4.0 DISCUSSION

### 4.1 DNA gyrase subunit A, GyrA

Interspecies MSA of the GyrA Ser81Leu mutant suggests that the variant is not prevalent among wild-type strains reported, with only one other selected strain (*A. baumannii* X2LSZ9) exhibiting a leucine residue, compared to the other strains exhibiting serine. The same is true for the intraspecies MSA between the ciprofloxacin-resistant PR07 and the ESKAPE pathogens, where only *S*.*aureus* exhibited a leucine residue, while the others exhibited serine, isoleucine, threonine. It is possible that the Ser81Leu is an uncommon mutation, especially among *A. baumannii*. This may be due to the resistant PR07 variant being exposed to ciprofloxacin in a controlled laboratory environment and lacking the dynamic interactions found in a host environment.

The Ser81Leu mutation in GyrA suggests a minor change in protein spatial occupancy, due to both serine and leucine being only three to four carbons in length (Betts & Russell, 2003). However, as leucine is Cβ branched (**Figure 2E**), it potentially increases the steric hindrance, altering the conformation of the DNA-GyrA complex. This may prevent ciprofloxacin binding by modifying the occupied molecular space (Betts & Russell, 2003). The mutation site is also positioned on the two GyrA monomers in such a way that they face each other, as well as the DNA molecule when complexed together (Gao et al., 2000).

Furthermore, the alteration from the polar serine to a non-polar leucine, suggests a potentially significant change in the biochemical activity of the 81th residue. The loss of the hydroxyl group (-OH) in serine would disable the formation of hydrogen bonds as well as any covalent bonds crucial for maintaining protein structure and function (Scheiner et al., 2001). This would influence either; i) the interaction between the GyrA monomers with DNA (Lin & Guo, 2019), or ii) the interaction between the two GyrA monomers themselves at the head dimer interface, which may in turn influence the conformation of the DNA-GyrAB complex. The change in polarity may influence the binding affinity of the negatively charged DNA; The loss of the slightly polar 81st residue could allow the DNA to more closely bind to the GyrA monomer (Betts & Russell, 2003). The loss of a hydroxyl group may also weaken the DNA-GyrA binding due to the inability to form a covalent bond (Mustaev et al., 2014)sure. This may alter the conformation of the DNA-GyrA complex, causing a reduction in the ciprofloxacin binding and increased resistance (Varughese et al., 2018).

In addition, DoGSiteScorer also determined the Ser81Leu mutation to be within proximity to a binding site in GyrA. Since the formation of the DNA-GyrAB complex involves the intertwining of the DNA molecule through the gate where the two GyrA monomers meet, it is possible that mutations within its proximity would alter binding activity (Couturier et al., 1998).

### 4.2 Efflux pump membrane transporter, MexB

Intraspecies MSA analysis of the meropenem-resistant PR07 variant revealed that the Ser181Leu mutation is not prevalent among the selected *A. baumannii* samples. However, small, aliphatic, non-polar amino acids are common, as exhibited by the four samples bearing alanine and glycine instead of the expected serine residue.

Interspecies MSA analysis also shows the same distribution of residues; glycine, alanine and serine among the samples. This indicates that these amino acids may be prevalent among wild-type strains for ESKAPE pathogens. In addition, the lack of leucine at position 181 of the MexB protein suggests that the mutation is unique to the PR07 variant, and may be a distinct outcome of the prolonged *in vitro* meropenem exposure.

Changes in amino acid structure and properties of the MexB Ser181Leu mutation are similar to the Ser81Leu mutation discussed in the GyrA variant. However, as the MexB protein functions as the embedded protein of the MexAB-OprM protein complex, the impact of the mutation is less clear. Since the mutation site is located on the surface of the internal tunnel of the MexB homotrimer (**Figure 4B**), it is possible that the structural changes incurred by the mutation could influence the passage of proteins through the efflux pump.

Similarly, with the GyrA mutation, the loss of a hydroxyl group (-OH) in the Ser181Leu variation may result in an altered efflux activity by reducing the protein binding energy (Betts & Russell, 2003). However, as efflux activity does not necessitate protein binding per se, it is unclear how this mutation would improve the antimicrobial resistance activity towards meropenem. While it is possible that some form of binding is necessary during entry into the OpRM-MexAB, the mutation site is located too far from the entry point at the lower portion of the MexB itself and may not play a significant role.

Potentially, the replacement of the electronegative serine with an aliphatic leucine would facilitate transversal across the MexAB-OprM internal passageway, since less binding activity is present. The presence of non-polar hydrophobic residues may improve hydrophobic sliding between proteins by switching from hydrogen bonds to van der Waals interactions (Foulkes-Murzycki et al., 2007). Furthermore, the increased hydrophobicity may slightly improve the stability of the protein (Gromiha et al., 1999).

### 4.3 Peptidoglycan D, FtsI

The intraspecies MSA of the imipenem-resistant PR07 strain with selected *A. baumannii* samples suggests that all three of the identified mutations; Pro508Leu, Ala515Val, and Ala579Thr are uncommon among the non-PR07 wildtype samples. It should be noted however, that the Ala579Thr was shown to be present in one of the samples (A0A334CLU8).

The FtsI protein is noted to be a part of a complex of proteins that play a crucial role in cell division, specifically the septa formation. Mutations in the membrane region of the FtsI protein have been previously associated with an increased rate of division to compensate for an increased cellular constriction brought on by sidA and didA mutations (Modell et al., 2014). However, there are as of yet no reports similar to the I45V variant discussed in the current study.

FtsI complexes with FtsW when forming the bacterial peptidoglycan, where the former plays the role of glycosyltransferase (GTase), and the latter functions as the transpeptidase (TPase). It was initially postulated that modifications in the FtsI would in some way alter the FtsWI complex interaction (Modell et al., 2014; Park et al., 2021). Literature suggests that FtsWI is complexed at the inner membrane, whereby FtsI protrudes into the intermembrane space (Rothfield & Justice, 1997). Furthermore, FtsI has been suggested to signal the activation of FtsW, likely through its pedestal region (Attaibi & den Blaauwen, 2022).

In the current study, the mutations identified were noted to be distal to where FtsI complexes with FtsW. All three polymorphisms occur at the surface of the FtsI monomer within the periplasmic space where the peptidoglycan synthesis occurs (**Figure 6A and Figure 6B**). The Pro508Leu variant occurs in a loop structure connecting an alpha-helix and beta-sheet, potentially impacting either the conformation of the catalytic region or modifying the transpeptidase activity.

The FtsI protein has also been reported to play an essential role in cell division, specifically the cross-linking of the bacterial peptidoglycan cell wall (Graham et al., 2021). It is possible that the mutations identified at the terminal region of the FtsI may affect the peptidoglycan synthesis process (Guzman et al., 1997; Wissel & Weiss, 2004). The exact effect is however unclear. Based on the findings of this study, the mutations occurring in the ftsI gene of PR07 (AbRI variant) upon exposure to imipenem are relevant, as the antibiotic targets the PBP-3 encoded by the gene. Analysis of the protein structure further confirms this as the mutations occur within the catalytic region of FtsI. Further study is needed to elucidate the exact impact of the amino acid alterations.

In regards to the mutation in the ftsI in the erythromycin variant, it is unclear as to what benefits the mutation may carry, as macrolides do not tend to interact with the peptidoglycan layer. At the time of this study, it can only be concluded that the ftsI mutation observed in the AbRE variant is likely a coincidental mutation with no added benefit to erythromycin resistance. However, further study is needed to confirm this.

### 4.4 ATP synthase subunit beta, AtpD

Both interspecies and intraspecies MSA analysis of the AtpD protein sequence revealed that the Ala166Val mutation is uncommon among the sequences acquired from NCBI. This suggests that the mutation may be unique to an *in vitro* setting.

Structural analysis of the Ala166Val mutation on the beta subunit of the ATP synthase suggests that there is no direct influence of the modification on the ATP synthesis activity, as it does not occur within any identified active sites (Ahmad et al., 2011; Cingolani & Duncan, 2011; Courbon & Rubinstein, 2022). Although there is very minimal change in the spatial occupancy between alanine and valine (**Figure 8D** and **Figure 8B**), the addition of a branched chain may alter the conformation of the protein to a degree (Ito et al., 2011; Guo & Rubinstein, 2022). This may affect the activity between the P-loop Clamp and the Active Site Clamp of the ATP synthase, potentially lowering cellular ATP production (Blum et al., 2012). Therefore, while the mutations were distal to the identified alpha-beta interface, where synthesis takes place, there is a possibility that the substrate-binding site of the Atp synthase subunit beta is affected by the Ala166Val mutation (Logan & Knight, 1993; Schmidt et al., 2001).

## 5.0 CONCLUSION

Overall, analyses of the protein alterations in the PR07 mutant strains revealed mostly uncommon polymorphisms in response to antibiotic exposure. Both MSA and protein modeling revealed novel mutations and structural changes that merit further study. While it is arguable that these mutations may be a result of culture bias, since the experiment was carried out under a controlled *in vitro* setting, the findings suggest mechanisms of adaptations that have yet to be observed in *A. baumannii*. Most notably, the mutations did little to modify the activity of binding pockets directly and instead suggested a change in protein-protein interactions. A more in-depth analysis of protein structure and dynamics would be needed to further extrapolate the impact of these mutations. Ultimately, the findings further emphasize the complexity of survival and adaptation mechanisms employed by *A. baumannii* when under antibiotics-induced selective pressure.

